# Working Memory Gates Visual Input to Primate Prefrontal Neurons

**DOI:** 10.1101/2020.11.09.375287

**Authors:** Behrad Noudoost, Kelsey L. Clark, Tirin Moore

## Abstract

Visually guided behavior relies on the integration of sensory input with information held in working memory. Yet it remains unclear how this is accomplished at the level of neural circuits. We studied the direct visual cortical inputs to neurons within a visuomotor area of prefrontal cortex in behaving monkeys. We show that the synaptic efficacy of visual cortical input to prefrontal cortex is gated by information held in working memory. Surprisingly, visual input to prefrontal neurons was found to target those with both visual and motor properties, rather than preferentially targeting other visual neurons. Furthermore, activity evoked from visual cortex was larger in magnitude, more synchronous, and more rapid, when monkeys remembered locations that matched the location of visual input. These results indicate that working memory directly influences the circuitry that transforms visual input into visually guided behavior.

## Introduction

Behavior is guided not only by sensory input, but also by information held in working memory (WM). In primates, visually guided eye movements are among the most frequently occurring sensorimotor transformations. Saccadic eye movements occur ∼4-5 times each second and require the integration of myriad visual features (e.g. motion and shape) into discrete movements that position visual targets onto the fovea. Furthermore, each movement decision reflects not only visual input, but also information held in WM, such as the behavioral relevance of particular objects and features (Bichot et al., 2005; Hollingworth et al., 2013; Hollingworth and Luck, 2009; Wolfe and Horowitz, 2004). How sensory input and WM are integrated in neural circuits to shape behavioral output remains unclear. Studies across multiple species have revealed evidence that WM functions are often associated with networks involved in sensorimotor transformations, including visual-saccadic transformations (Curtis and D’Esposito, 2006; Gnadt and Andersen, 1988; Guo et al., 2014; Knudsen et al., 1995; Knudsen and Knudsen, 1996; Kojima et al., 1996). In these networks, neurons, individually or collectively, often exhibit persistent signaling of information needed to successfully carry out subsequent behaviors. The prevalence of WM-related activity in sensorimotor networks suggests that this may be where WM exerts its influence on sensorimotor transformations. However, the exact mechanism and specific neural circuitry by which WM influences visually guided behaviors are still unknown.

Within primate neocortex, the output of feature-selective neurons in visual cortical areas converges retinotopically onto neurons in the frontal eye field (FEF)(Schall et al., 1995), the prefrontal area mostly directly involved in the control of saccades (Robinson and Fuchs, 1969; Schiller et al., 1979). Neurons within the FEF exhibit functional properties spanning the visual-motor spectrum, and also include a substantial portion of neurons with persistent, WM-related activity (Bruce and Goldberg, 1985; Lawrence et al., 2005). These characteristics make the FEF a likely place to observe an influence of WM on incoming visual signals, particularly given that the FEF transmits a predominantly WM signal to visual cortex (Merrikhi et al., 2017). Although much is understood about the role of FEF neurons in the control of visually guided saccades (Schiller et al., 1979; Schlag-Rey et al., 1992; Tehovnik et al., 2000), and in the control of visual spatial attention (Bahmani et al., 2019; Gregoriou et al., 2009; Moore and Fallah, 2001), very little is known about how visual, motor, and memory signals are combined within the FEF. Models of FEF microcircuitry generally predict that visual cortical inputs synapse predominantly onto purely visual FEF neurons (e.g. Heinzle et al., 2007), yet even this is not known. Furthermore, it is also not known how those visual inputs interact with the current content of WM.

We examined the influence of WM on the efficacy of visual cortical input to the FEF in behaving monkeys. First, we identified FEF neurons with direct input from visual cortex using orthodromic stimulation from extrastriate area V4. Despite the common assumption of visual inputs synapsing onto purely visual FEF neurons, our results revealed that visual cortical input to the FEF instead preferentially targets neurons with both visual and motor properties. Next, we measured the effect of spatial WM on orthodromic activation of FEF neurons and found that the synaptic efficacy of visual inputs was enhanced by the memory of spatially corresponding locations. Specifically, the activity evoked in the FEF from visual cortex was larger in magnitude, more synchronous, and more rapid when V4 input matched the location held in WM. These results demonstrate how the content of WM can influence visuomotor transformations in the primate brain.

## Results

We measured the influence of WM on the efficacy of visual cortical inputs to prefrontal cortex in behaving monkeys. Monkeys performed a spatial WM task classically used to characterize FEF neuronal properties (Fig. 1a)(Bruce and Goldberg, 1985; Lawrence et al., 2005). Figure 1b shows the activity of an example FEF neuron when a monkey remembered a location either inside or outside of the response field (RF; *In* and *Out* conditions, respectively). The neuron responded strongly to a visual cue appearing in the RF and exhibited elevated activity in the delay period when the remembered location coincided with the RF. Prior to saccades to the RF, the neuron also exhibited a burst of motor activity. In primates, direct visual cortical input to prefrontal cortex arrives primarily in the FEF (Markov et al., 2014a; Ungerleider et al., 2008). We orthodromically activated FEF neurons from retinotopically corresponding sites in extrastriate area V4 (Fig. 2a)(Methods). Of 311 single FEF neurons recorded, spikes were reliably elicited by V4 stimulation in 115. Latencies of evoked spikes were bimodally distributed (Hartigan’s dip test, p<10^−40^), with most neurons having latencies <10 ms (n=96, latency=6.53±0.67 ms), consistent with monosynaptic transmission (Fig. 2b)(Gregoriou et al., 2009; Nowak LG, Bullier J., 1997). We focused our analyses on these *visual-recipient* neurons. A smaller population of neurons was activated at longer latencies (n=19, latency=12.73±1.40 ms). Visual-recipient neurons exhibited a tri-phasic pattern of evoked activity following orthodromic stimulation (Fig. 2c, S1), similar to previous studies employing orthodromic stimulation in primate neocortex (Matsunami and Hamada, 1984).

**Figure 1.**
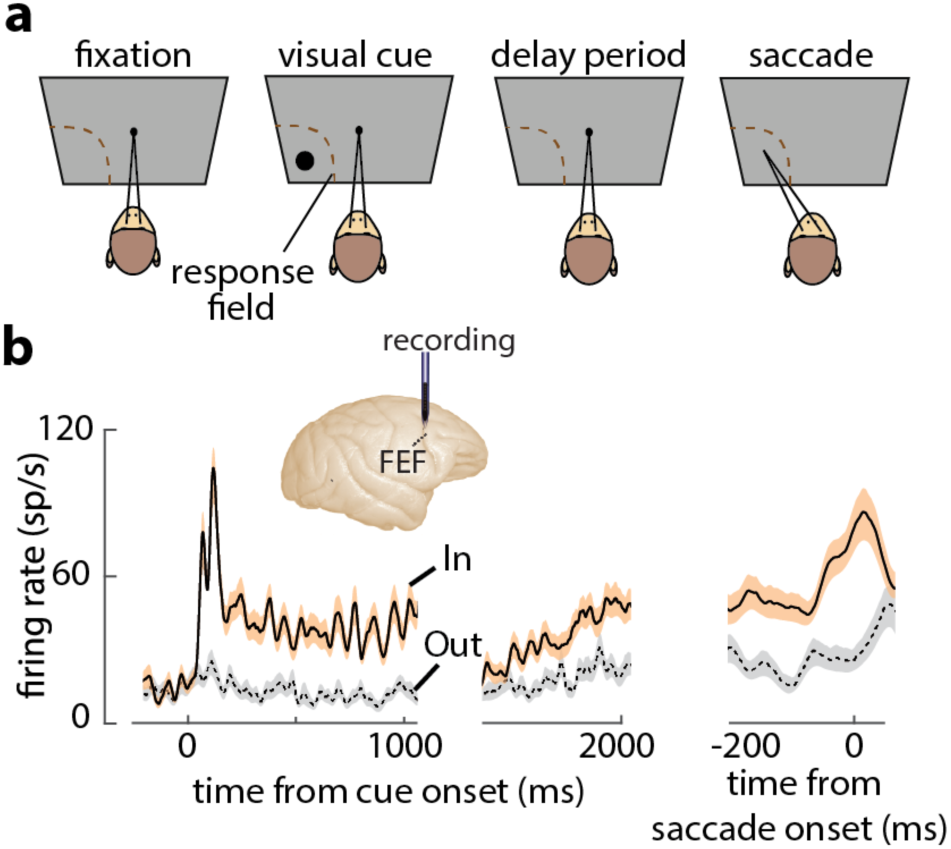
FEF responses during the memory guided saccade (MGS) task. A) Schematic of the MGS task. Monkey fixates, and a visual cue is presented (inside or outside the neuronal RF). The monkey maintains fixation throughout a delay period, and upon removal of the fixation point, saccades to the remembered location to receive a reward. B) Response of an example FEF neuron during the MGS task; this neuron shows visual, memory, and motor activity. Responses are shown for cues inside (In, orange) or outside (Out, gray) the RF, aligned to cue onset (left, middle panels) or the saccade (right panel). Traces show mean ± SEM.

**Figure 2.**
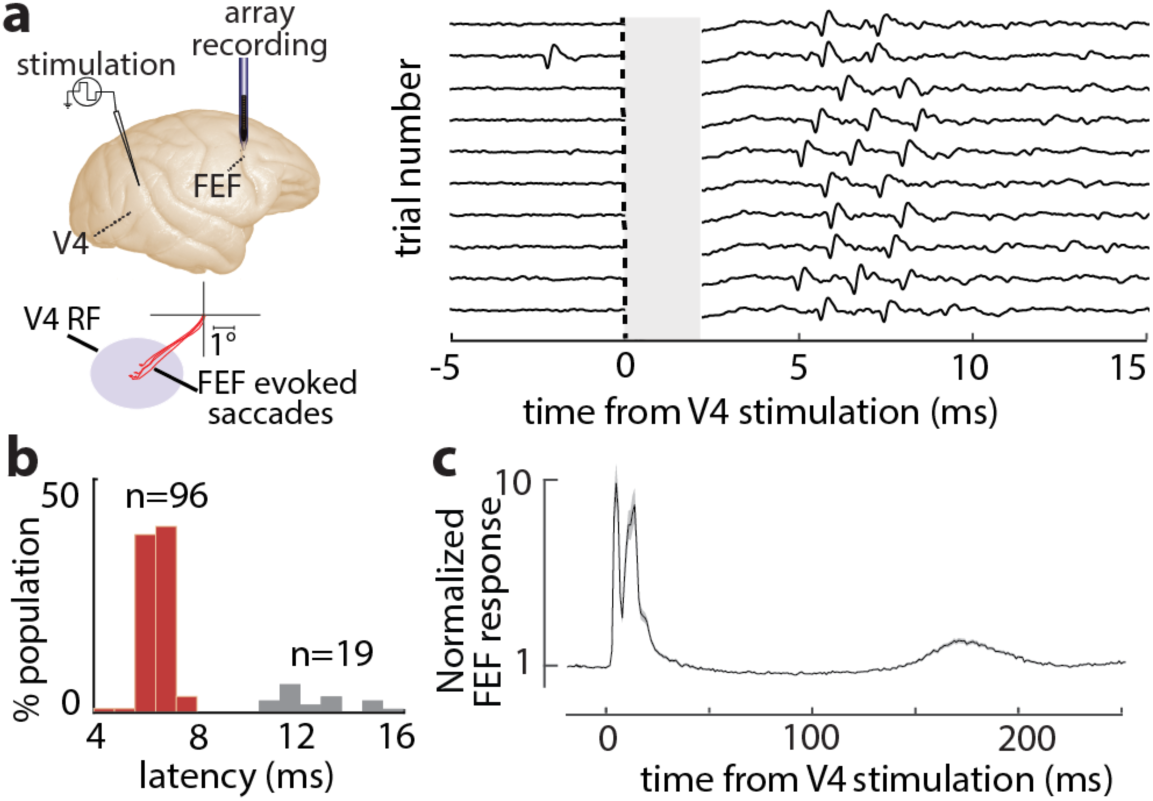
Orthodromic activation of FEF neurons from visual cortex. A) FEF neurons were orthodromically activated by electrical stimulation of retinotopically corresponding V4 sites (left). Right plot shows evoked spikes from an FEF neuron across 10 trials (stimulation artifact period is shown in gray). B) Distribution of stimulation-evoked spike latencies for 115 orthodromically activated FEF neurons. C) Average normalized stimulation-evoked activity of the 96 visual-recipient FEF neurons over time.

### Characterizing response properties of visual-recipient neurons

We tested whether the properties of visual-recipient neurons differed from those not activated by orthodromic stimulation (*non-activated* neurons)(n=196). We measured the activity of FEF neurons while monkeys performed a spatial WM task that temporally dissociates visual, memory, and motor components of neuronal responses (Methods). Figure 3a shows the activity of an example visual-recipient FEF neuron when the monkey remembered a location either inside or outside of the RF. This neuron responded strongly to a visual cue appearing in the RF but exhibited minimal activity in the delay period, and was not selective for the remembered location. Prior to saccades to the RF, the neuron exhibited a burst of motor activity. Thus, this neuron exhibited visual and motor, not memory-related, activity. We compared the proportions of neurons with significant visual, memory, and motor activity between the visual-recipient and non-activated neuronal populations. Each component of activity was measured as the selectivity between In and Out conditions in the corresponding behavioral epoch. We found that visual-recipient FEF neurons exhibited greater proportions of visual (χ^2^=9.42, p=0.002) and motor activity (χ^2^=10.71, p=0.001) than non-activated neurons. However, the proportion of neurons with memory activity did not differ between the two populations (χ^2^=0.99, p=0.318)(Fig. 3b-left). We further compared the proportions of neurons exhibiting both visual and motor activity (visuomotor), or only visual or motor activity, between the visual-recipient and non-activated populations. Overall, the relative proportions of visual, visuomotor, and motor neurons differed between the visual-recipient and non-activated populations (χ^2^=6.89, p=0.0319), with a higher prevalence of visuomotor neurons among the visual-recipient population (66% vs. 44%, χ^2^=11.34, p<10^−3^), and a lower proportion of purely visual (19% vs. 31%, χ^2=^4.39, p=0.036) and purely motor neurons (15% vs. 26%, χ^2^=4.23, p=0.039)(Fig. 3b-right). Thus, the increased prevalence of visual and motor activity among the visual-recipient neurons reflected a larger proportion of visuomotor neurons.

**Figure 3.**
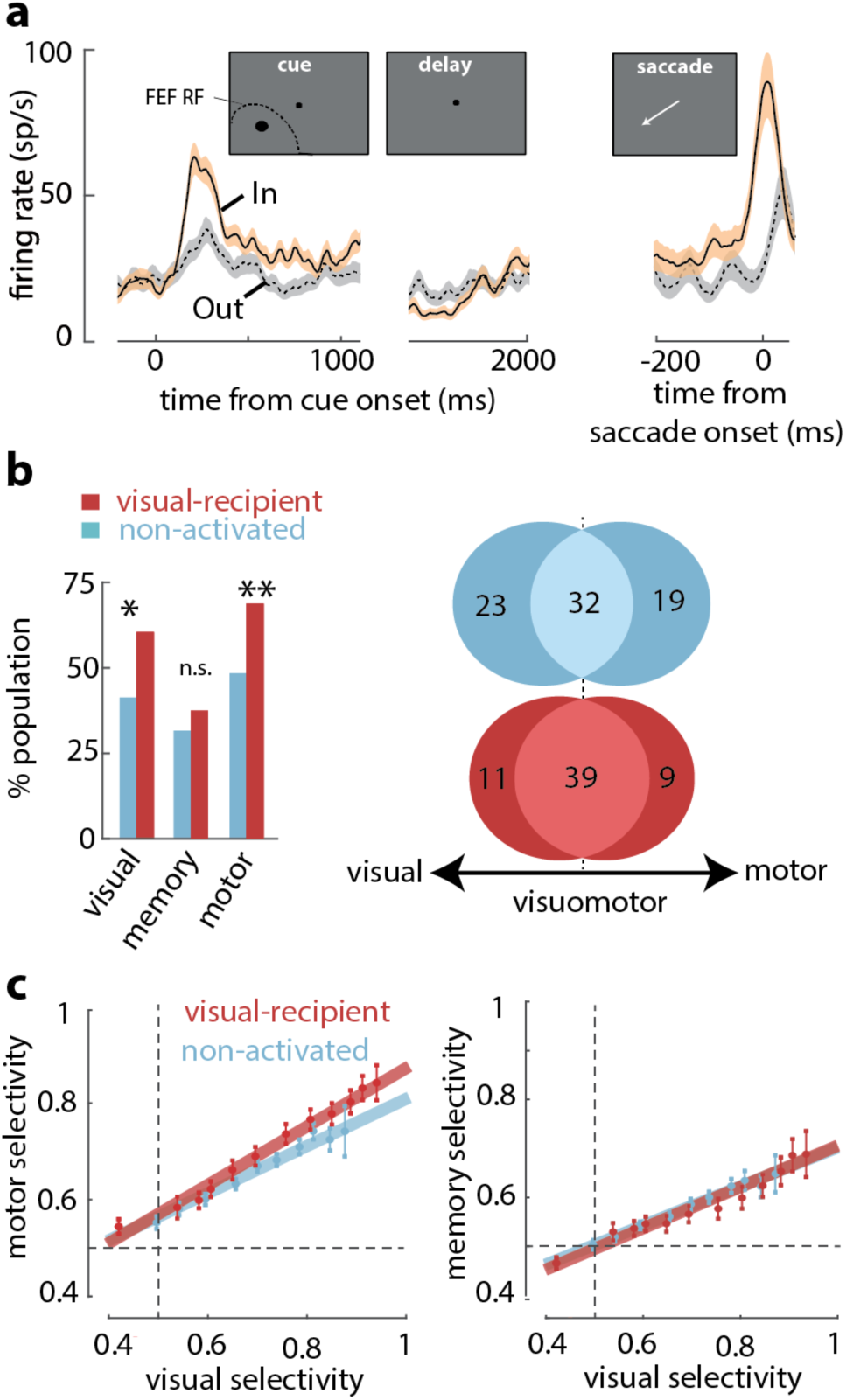
Overrepresentation of visuomotor activity in visual-recipient FEF neurons. A) Activity of an example visual-recipient FEF neuron when the cue appeared In (peach) or Out (gray) of the RF. Plots show firing rate aligned to the onset of the visual cue (left), offset of the visual cue (middle), and to saccade onset (right). B) Left: Percent of the population exhibiting visual, memory, and motor activity for visual-recipient (cardinal) and non-activated (turquoise) FEF neurons. * and ** denote p<0.05 and p<0.001 respectively and n.s. denotes p>0.05. Right: Venn diagrams show number of neurons with visual, motor, or visuo-motor activity for visual-recipient (top) and non-activated (bottom) FEF neurons. C) Motor selectivity (left) and memory selectivity (right) as a function of visual selectivity for visual-recipient and non-activated FEF neurons.

We considered that the larger motor signals among visuo-recipient neurons could have resulted from differences in the alignment of the cue stimulus with the centers of FEF visual and movement fields, as they can be significantly misaligned (Bruce and Goldberg, 1985; Schafer and Moore, 2011). Thus, we measured the magnitude of motor activity (selectivity to In vs. Out) across varying amounts of visual in the two populations of neurons (Methods) (Fig. 3c-left). This comparison revealed that for a given level of visual activity, visual-recipient neurons exhibited a larger component of motor activity than non-activated neurons (ANCOVA, F=10.15, p=0.002). In contrast, a corresponding comparison of memory and visual activity in the two populations revealed no differences (ANCOVA, F=0.23, p=0.631)(Fig. 3c-right). Thus, whereas memory signals among visual-recipient FEF neurons were equal to those of non-activated neurons, motor signals were significantly overrepresented (See table S1 and Fig. S2).

### WM alters efficacy of V4 input to FEF

In contrast to the disproportionate prevalence of motor signals among neurons receiving input from visual cortex, reciprocal projections of the FEF to visual cortex originate disproportionately from neurons with memory-related activity (Merrikhi et al., 2017). This implicates the FEF as a possible source of the observed memory-dependent modulation in visual cortex (Bahmani et al., 2019, 2018). It also suggests that memory-related signals may interact with the efficacy of visual inputs arriving in prefrontal cortex. To test this possibility, we measured the effects of engaging WM on the efficacy of orthodromic activation of FEF neurons from V4. Previous studies show that the efficacy of orthodromic activation of primate V1 neurons from the thalamus (Briggs et al., 2013), and of extrastriate cortex from primary visual cortex (Ruff and Cohen, 2016), are both altered during spatial attention. Using a similar approach, we compared the activity evoked from visual cortex by orthodromic stimulation during periods when monkeys remembered different cue locations.

We examined the influence of WM across the full population of 96 visual-recipient neurons. On average, the proportion of stimulation-evoked spikes increased by 19% when monkeys remembered locations inside the RF, compared to outside (Spike count_In_=0.22±0.005; Spike count_Out_=0.18±0.004; p<10^−6^)(Fig. S3). For each neuron, we measured the evoked response magnitude to quantify the efficacy of stimulation (see Methods). The magnitude of evoked responses was significantly greater when monkeys remembered cue locations inside the RF, compared to outside (Response magnitude_Out_=0.68±0.03; Response magnitude_In_=0.77±0.04; p<10^−10^) (Fig. 4a). Thus, the efficacy of visual input to the FEF depended on the content of WM. In addition, we found that the latency of evoked spikes was slightly reduced when monkeys remembered locations inside, compared to outside, the RF (Latency_In_=7.88, Outside_RF_=8.04; p<10^−10^)(Fig. 4b). Figure 4c shows the spikes evoked from two example FEF neurons in response to V4 stimulation during the memory period. Evoked spikes from one neuron increased by ∼30% during memory of locations inside, compared to outside the RF. The number of evoked spikes from a second, simultaneously recorded, neuron was similar in the two RF conditions, but spike onset appeared more rapid during the In condition, consistent with the latency effects. As a consequence, when combined, the evoked spikes of the two neurons were more synchronous during memory of locations inside of the RF (Fig. 4c)(Methods). We compared the probability of joint spiking in all simultaneously recorded pairs of visual-recipient neurons (n=509) across the different memory locations. Overall, we found that the probability of joint spiking, when controlled for firing rate (see Methods), increased by nearly 60% during memory of locations inside, compared to outside the RF (Prob_In_=0.103±0.002, Prob_Out_=0.065±0.001, p<10^−63^)(Fig. 4d). Thus, activity evoked in the FEF from visual cortex was larger, more synchronous, and more rapid when monkeys engaged WM at RF locations.

**Figure 4.**
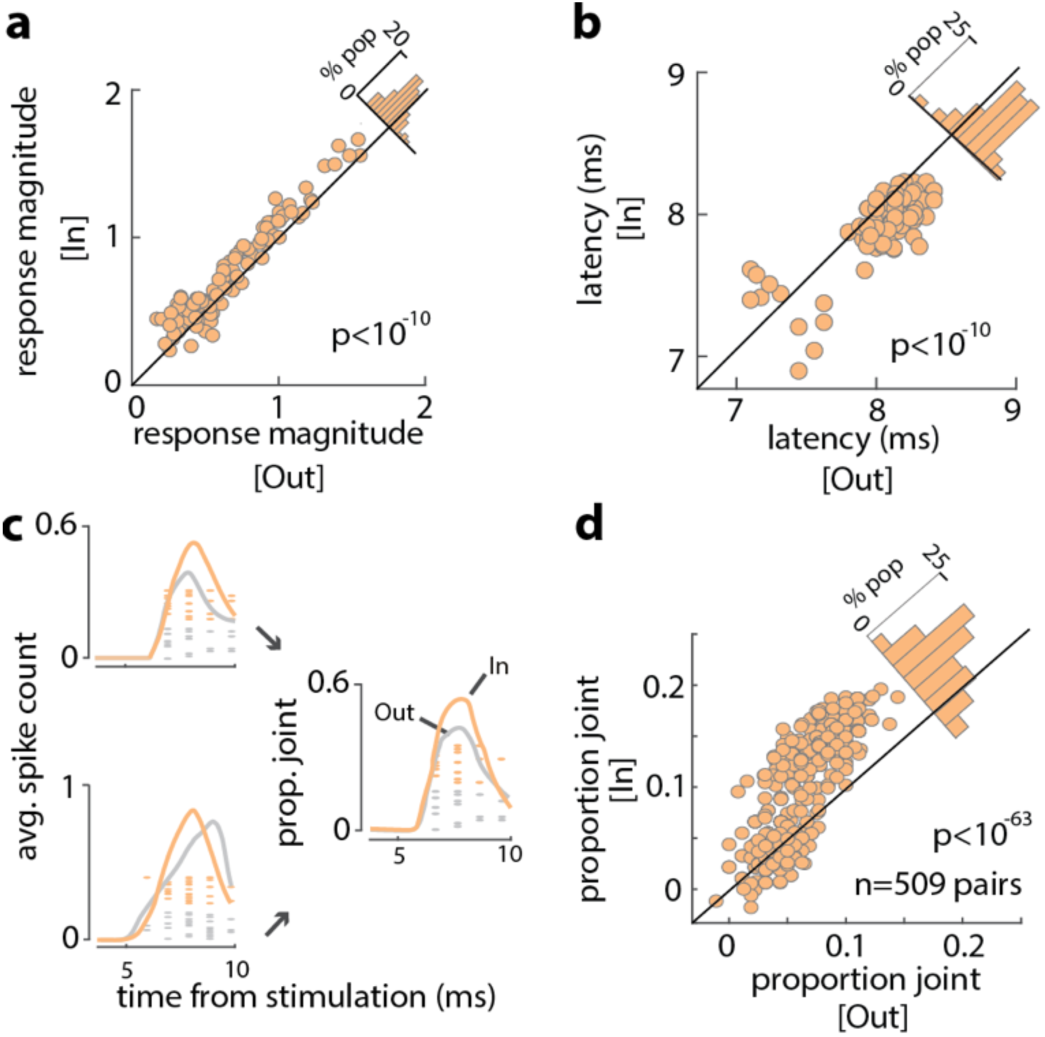
Increased efficacy of visual input to the FEF during WM. A) Magnitude of orthodromically evoked FEF responses, and B) latency of orthodromically evoked spikes during memory of locations inside and outside of the RF for all visual-recipient FEF neurons. C) Left: mean spike counts and raster plots following V4 stimulation for 2 example neurons during memory of locations inside (peach) and outside (gray) of the RF. Right: Proportion of joint spikes in the two example neurons during the two memory conditions. D) Mean proportion of joint spikes for all pairs of visual-recipient FEF neurons during the two memory conditions.

## Discussion

We found that visual inputs to the FEF disproportionately drove neurons with both visual and motor properties. Rather than exhibiting purely visual properties, as might be expected (Sato and Schall, 2003), visual-recipient FEF neurons also signaled the direction of impending eye movements. Although surprising, this result seems consistent with the pattern of visual cortical connections with the FEF (Barone et al., 2000; Markov et al., 2014b). V4 inputs terminate in all layers of the FEF (Ungerleider et al., 2008), thus potentially distributing inputs across different functional classes of neurons. The bias in those inputs toward visuomotor neurons indicates that rather than being integrated at a subsequent processing stage within the FEF, as proposed by models of FEF microcircuitry (Brown et al., 2004; Heinzle et al., 2007), sensory and motor signals are combined at the interface between visual and prefrontal cortex.

More importantly, we found that the engagement of WM at RF locations increased the efficacy of visual input to the FEF. When monkeys remembered RF locations, activity evoked in the FEF from visual cortex was larger in magnitude, more synchronous, and more rapid. Increases in synaptic efficacy have previously been observed in thalamocortical (Briggs et al., 2013) and corticocortical (Ruff and Cohen, 2016) connections during the deployment of attention. Evidence from multiple species suggests that top-down attention arises from biases in the selection of sensory input based on the content of WM (Desimone and Duncan, 1995; Knudsen, 2007; Miller and Cohen, 2001). A recent study indicates that FEF neurons with WM-related activity disproportionately provide input to neurons in area V4 (Merrikhi et al., 2017), and thus underlie the FEF’s contribution to the modulation of visual activity classically observed during spatial attention (Ekstrom et al., 2009; Gregoriou et al., 2014; Moore and Armstrong, 2003). In contrast to visual cortical projections from the FEF, inputs to the FEF did not preferentially target memory-related neurons, but instead neurons with visuomotor properties. This suggests that interactions between the FEF and visual cortex are not strictly recurrent (Knudsen, 2007; Noudoost et al., 2014), and that memory activity within the FEF, rather than reinforcing its own content, may instead facilitate the transformation of visual inputs into motor commands. Combined, our results suggest a basis for the well-document interdependence of attention, WM and gaze control (Ikkai and Curtis, 2011; Jonikaitis and Moore, 2019), which at the circuit-level, remains an important puzzle to solve.

## Methods

Two adult male rhesus monkeys (*Macaca mulatta*) were used in this study. All experimental procedures were in accordance with National Institutes of Health *Guide for the Care and Use of Laboratory Animals*, the Society for Neuroscience Guidelines and Policies, and Stanford University Animal Care and Use Committee.

### General and surgical procedures

Each animal was surgically implanted with a head post, a scleral eye coil, and two recording chambers. Two craniotomies were performed on each animal, allowing access to dorsal V4, on the prelunate gyrus, and FEF, on the anterior bank of the arcuate sulcus. Eye position monitoring was performed via the scleral search coil and was digitized at 500 Hz (CNC Engineering). Eye monitoring, stimulus presentation, data acquisition, and behavioral monitoring were controlled by the CORTEX system. Visual stimuli presented to estimate V4 RFs were 1.2−1.9° × 0.2−0.4° bar stimuli appearing at four possible orientations (0, 45, 90, and 135°). All stimuli were presented on a 29° × 39° (22″) colorimetrically calibrated CRT monitor (Mitsubishi Diamond Pro 2070SB-BK) with medium short persistence phosphors (refresh rate 77 Hz).

### Neurophysiological recording and data acquisition

Neurophysiological recordings of single neurons in awake monkeys were made through two surgically implanted cylindrical titanium chambers (20 mm diameter) overlaying the prelunate gyrus (V4) and the pre-arcuate gyrus (FEF). Electrodes were lowered into the cortex using a hydraulic microdrive (Narishige). Neural activity was recorded extracellularly with varnish-coated tungsten microelectrodes (FHC) of 0.2–1.0 MΩ impedance (measured at 1 kHz) in V4, and via linear electrode array (Plexon, v-probe) in FEF. Extracellular waveforms were digitized and classified as single neurons using both template matching and window-discrimination techniques (FHC, Plexon). Area V4 was identified based on stereotaxic location, position relative to nearby sulci, patterns of grey and white matter, and response properties of units encountered; the FEF was identified based on these factors and the ability to evoke fixed-vector eye movements with low-current electrical stimulation. Prior to beginning data collection, the location of FEF and V4 within the recording chambers was established via single-electrode exploration.

#### Eye Calibration

Each day began by calibrating the eye position; once the electrode was positioned in the FEF, the same task was used with stimulation to verify that the electrode was in FEF and estimate the RF center. The fixation point, a ∼1 degree of visual angle (d.v.a.) white circle, appeared in the center of the screen, and the monkey maintained fixation within a ±1.5 d.v.a. window for 1.5 s. For eye calibration, no stimulation was delivered and the fixation point could appear either centrally or offset by 10 d.v.a. in the vertical or horizontal axis.

#### Achieving FEF-V4 overlap

In each recording session, we first localized sites within the FEF and V4 where neurons exhibited retinotopically corresponding representations, meaning that V4 RFs overlapped with the end point of saccade vectors evoked by FEF microstimulation(Merrikhi et al., 2017; Moore and Armstrong, 2003). Preliminary RF mapping in V4 was conducted while the monkey fixated within a ±1.5 d.v.a. window around the central fixation point, while ∼2.5 × 4 d.v.a. white bars swept in eight directions (four orientations) across the approximate location of the neuron’s RF. Responses from the recording site were monitored audibly and visually by the experimenter, and the approximate boundaries of the RF were noted for the positioning of stimuli in subsequent behavioral tasks. To establish that the electrode was positioned within the FEF and to estimate the FEF RF location, microstimulation was delivered randomly on 50% of trials while the animal performed a passive fixation task. Microstimulation consisted of trains (50–100 ms) of biphasic current pulses (≤50 μA; 250 Hz; 0.25 ms duration). On no-stimulation trials, the monkey was rewarded for maintaining fixation; on stimulation trials, the monkey was rewarded whether fixation was maintained or not following microstimulation. The ability to evoke saccades with low stimulation currents (≤50 μA) confirmed that the electrode was in the FEF; the end point of the stimulation-evoked saccades provided an estimate of the RF center for the FEF site.

#### Memory-guided saccade (MGS) task

The FEF visual, motor, and delay activity were characterized in an MGS task. Monkeys fixated within a ±1.5 d.v.a. window around the central fixation point. After 1 s of fixation, a 1.35 d.v.a. square cue was presented and remained onscreen for 1 s. The animal then remembered the cue location while maintaining fixation for 1 s (delay period) before the central fixation point was removed. The animal then had 500 ms to shift its gaze to a ±4 d.v.a. window around the previous cue location and remain fixating there for 200 ms to receive a reward. This task was performed with two potential cue locations, located at 0° and 180° relative to the estimated RF center.

#### Electrical stimulation

During the MGS task described above, electrical stimulation was delivered to V4 during the fixation, visual, delay, or saccade period on 50% of trials (on the other 50% of trials there was no stimulation). For identifying antidromically and orthodromically activated FEF neurons (see below), and evaluating stimulation efficacy, electrical stimulation consisted of a single biphasic current pulse (600–1,000 μA; 0.25 ms duration, positive phase first). Stimulation times were 500 ms after initiating fixation (fixation), 500 ms after visual cue onset (visual), 500 ms after cue offset (delay), or 150 ms after the go cue (saccade).

### Statistical analysis

#### Latency of stimulation evoked spikes

The probability of firing in each 1 ms bin following V4 stimulation was measured for stimulation trials and compared to the probability of firing in a time-matched window from non-stimulation trials. The first bin in which the probability of firing was significantly greater for stimulation trials was designated the latency of stimulation-evoked spikes. Hartigan’s dip test was used to test the bimodality of the latency distribution (Fig. 2B).

#### Identifying orthodromically activated neurons

Electrical stimulation of V4 evoked spikes in FEF via both orthodromic and antidromic stimulation. Antidromically-evoked spikes (in V4-projecting FEF neurons) were of short latency and confirmed via the collision test (using stimulation data collected during the MGS task described above). This test identifies antidromically activated neurons: when V4 stimulation was delivered within a few milliseconds of a spontaneously generated spike from a recorded FEF neuron, spikes artificially evoked from that neuron by V4 microstimulation were eliminated. Orthodromically activated neurons will still have an evoked spike in this period following a spontaneously generated spike. The characteristics of FEF neurons antidromically activated by V4 stimulation have been reported elsewhere(Merrikhi et al., 2017).

#### Assessment of stimulation-evoked activity

All responses are measured within the 5-9 ms post-stimulation period, to stay consistent with the latency of visual-recipient neurons as shown in figure 2b. To focus on stimulation-evoked spikes, rather than spontaneous spiking activity, all measures are adjusted by subtracting the same measure (firing rate, spike count, or probability) observed during the same time period on non-stimulated trials. Figure 4 shows adjusted values, after the subtraction of the same measure during the non-stimulated trials. The evoked response magnitude (Fig. 4a) was calculated based on the log ratio of spike counts before vs. after stimulation, and subtracting the same measure during non-stimulations trials: log_10_(resp after/resp before)_STIM_ - log_10_(resp after/resp before)_NONSTIM_. Probability of joint spiking (Fig. 4d) was defined as simultaneously recorded neurons firing within 1ms of each other, and was averaged from 5-9ms post-stimulation. Greater firing rates will increase the probability of joint spiking. To have a firing-rate independent measure of joint activity between the two neurons for each condition, we removed the temporal relationship between the two neurons (ie, shuffled the trial pairings) and subtracted the measured joint probability from that measured when the two neurons were simultaneously recorded. Similar to other measures this rate-matched value in non-stimulated trails was also subtracted from the stimulated trials. Thus, the reported joint probability is controlled for enhanced firing rate due to both stimulation and WM, and measures only the change in synchronous firing independent of firing rate.

#### Characterizing FEF response properties

The visual, motor and delay period activity of FEF neurons were measured using the MGS task described above. The visual period included activity 100–1,000 ms after stimulus onset. Delay period activity was measured from 300 to 1,000 ms after stimulus offset. Motor activity was quantified in the perisaccadic window from 75 ms before to 25 ms after the saccade onset. These time windows were also used to quantify the different types of activity using an ROC selectivity analysis described below. When determining whether a neuron had significant visual or delay activity, activity in the visual and delay periods of the In condition was compared to the activity of the same neuron during fixation (300 ms before stimulus onset), using the Wilcoxon sign-rank test (*P*<0.05). When determining whether a neuron had significant motor activity, saccade-aligned activity in the In condition was compared to saccade-aligned activity earlier in the trial (450–250 ms before saccade onset), using the sign-rank test (*P*<0.05).

The strength of visual, memory and motor activity was measured as the selectivity of neurons to the In and Out conditions during the visual, delay and motor epochs, respectively, and was quantified using the ROC method. This method compared the distributions of firing rates for trials in which the memory cue appeared inside versus outside the neuron’s RF(Green and Swets, 1966). The areas under ROC curves were used as a measure of selectivity for cue location, and were calculated as in previous studies(Armstrong and Moore, 2007; Britten et al., 1992). Specifically, we computed the average firing rate in the visual, delay and saccade windows defined above, for In and Out trials. We then computed the probability that the firing rate in each stimulus condition exceeded a criterion. The criterion was incremented from 0 to the maximum firing rate, and the probability of exceeding each criterion was computed. Thus, a single point on the ROC curve is produced for each increment in the criterion, and the entire ROC curve is generated from all of the criteria. The area under the ROC curve is a normalized measure of the separation between the two firing rate distributions obtained when the WM cue appeared inside versus outside the neuronal RF and provides a measure of how well the neuronal response discriminates between the two conditions.

## Acknowledgements

This work was supported by National Institutes of Health grants R01EY02694, R01NS113073, and R01MH121435 to BN, an Unrestricted Grant from Research to Prevent Blindness, Inc. to Moran Eye Center, University of Utah, and R01EY014924 and Howard Hughes Medical Institute grants to TM.

## Author Contributions

Experiments were designed by BN and TM and performed and analyzed by BN and KLC. TM, KLC and BN prepared the manuscript.

