## Supplementary Information for "Working Memory Gates Visual Input to Primate Prefrontal Neurons"

**Supplemental Information**

**Timecourse of FEF response enhancement and suppression following V4 stimulation.**

Visual-recipient neurons were defined based on a short-latency increase in spiking probability following V4 stimulation. However, in addition to this short-latency response, a consistent response dynamic was observed across the population of visual-recipient neurons with an initial burst of spikes (3.71±1.60 to 29.99±6.73 ms), followed by a period of suppression (57.06±19.84 to 102.10±37.40 ms), and then a later period of enhancement (162.52±11.24 to 193.37±14.15 ms), relative to baseline activity (Fig. 1c and supplementary Fig. S1). Figure S1A shows the proportion of the visual-recipient population that is enhanced or suppressed over time, along with the average mean-normalized response of the population. Note that the average increase in firing rate within ~30ms of stimulation was much larger than the subsequent increase in firing rate ~150-200ms following stimulation, as one would expect for direct stimulation of monosynaptic inputs. Figure S1B shows the time windows in which the response of each of the 96 visual-recipient neurons was significantly enhanced or suppressed; timing of the stimulation-induced changes in firing rate was fairly consistent across neurons.

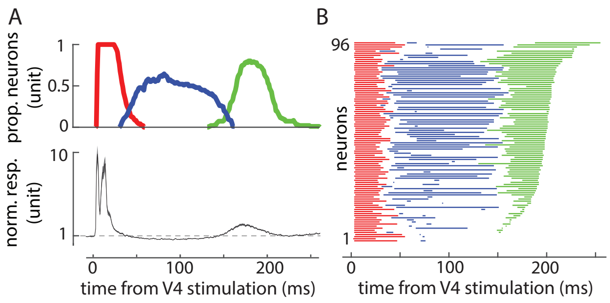


**Supplemental Figure 1)** Enhancement and suppression of visual-recipient FEF neurons' activity following electrical stimulation of V4. A) Top: proportion of the visual-recipient FEF population showing significantly elevated firing rates (red and green), or significantly suppressed firing rates (blue), over time relative to V4 stimulation. Firing rates are compared to a time-matched window on non-stimulation trials. Bottom: normalized response of the visual-recipient population following stimulation (mean±SE). The tri-phasic pattern of enhancement,-suppression-enhancement is a common characteristic of visual-recipient neurons. Significance of change in firing rate for each of the 96 visual-recipient FEF neurons is shown in B. Significance of stimulation-induced change in each neuron's response is plotted in a row, over time relative to V4 stimulation. Red and green indicate a significant increase in firing rate. Blue indicates a significant decrease in firing rate. White: no significant change. (Neurons are ordered based on the total number of time bins reaching significance.)

**Comparison of visual, memory, and motor selectivity in the visual-recipient, slow-input, and non-activated FEF populations**.

In order to compare the selectivity seen in the visual-recipient and non-activated populations (n = 96 and 196 respectively) with each other and with the smaller sample of slow-input population (n=19), the visual-recipient and non-activated populations were subsampled 1000 times. Sub-samples of 19 neurons were drawn from the population of visual-recipient and non-activated neurons in order to compare the magnitude of selectivity in these subsamples with each other and the slow-input group. Figure S2 shows the distribution of the visual, memory, and motor selectivity of these subsamples in visual-recipient and non-activated sub-samples and the single value for the slow-input group (black line). Visual-recipient neurons displayed much greater visual and motor selectivity than both the slow-input or non-activated populations ($p<10^{-3}$). Table 1 shows the average visual, memory, and motor selectivity values for the various FEF populations, and the significance of their statistical comparisons between them.

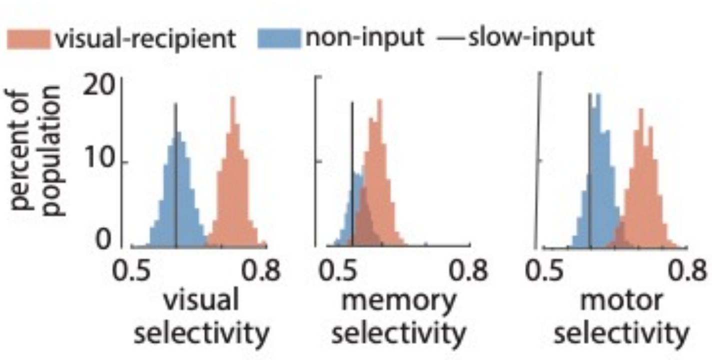


**Supplemental Figure 2)** Distribution of visual, memory, and motor selectivity for random subsamples of non-input (blue) and visual-recipient (red) FEF neurons. Black line indicates median selectivity of slow-input FEF neurons (n= 19). To better compare the visual-recipient and non-activated populations to the much smaller slow-input population, the histograms show distributions of average selectivity of 1,000 19-neuron ensembles, each selected randomly from the visual-recipient or non-activated populations (n = 96 and 196, respectively).

<table>
<tr><th></th><th>visual selectivity</th><th colspan="3">comparison</th><th>memory selectivity</th><th colspan="3">comparison</th><th>motor selectivity</th><th colspan="3">comparison</th></tr>
<tr><td>Non-activated (n=196)</td><td>0.603± 0.006</td><td rowspan="2">$p<10^{-3}$</td><td></td><td rowspan="3">$F=22.69, p<10^{-3}$</td><td>0.545± 0.007</td><td rowspan="2">p=0.023</td><td></td><td rowspan="3">F=4.69, p=0.010</td><td>0.619± 0.008</td><td rowspan="2">$p<10^{-3}$</td><td></td><td rowspan="3">$F=14.07, p<10^{-3}$</td></tr>
<tr><td>Visual-recipient (n=96)</td><td>0.720± 0.014</td><td rowspan="2">$p<10^{-3}$</td><td>0.586± 0.013</td><td rowspan="2">p=0.071</td><td>0.702± 0.015</td><td rowspan="2">p=0.003</td></tr>
<tr><td>Slow-input (n=19)</td><td>0.560± 0.018</td><td></td><td>0.530± 0.020</td><td></td><td>0.599± 0.024</td><td></td></tr>
<tr><td>all (n=311)</td><td>0.639± 0.009</td><td colspan="3"></td><td>0.557± 0.007</td><td colspan="3"></td><td>0.643± 0.008</td><td colspan="3"></td></tr>
</table>

**Supplemental Table 1)** Statistical comparisons of the visual, memory, and motor selectivity for non-activated, slow-input, and visual-recipient FEF neurons. Selectivity calculated based on the area under the ROC curve for In vs. Out conditions; standard error calculated across neurons. P-values for comparisons between two groups of neurons calculated using the Wilcoxon signed-rank test; comparisons across all three groups calculated using the ANOVA.

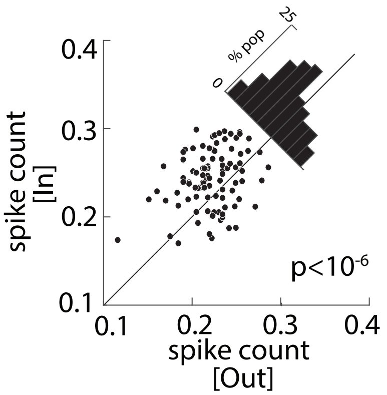


**Supplemental Figure 3)** Average spike count as a measure of stimulation efficacy across the population of visual-recipient FEF neurons, for memory Out (x-axis) vs. In (y-axis); diagonal histograms show differences. Remembering the RF location increased the average spike count within the 5-9 ms after stimulation by 19% for the population of 96 of visual-recipient FEF neurons (In = 0.22±0.005; Out = 0.18±0.004; P $<10^{-6}$).
